# Two conformations of the Tom20 preprotein receptor in the TOM holo complex

**DOI:** 10.1101/2023.01.26.525638

**Authors:** Pamela Ornelas, Thomas Bausewein, Janosch Martin, Nina Morgner, Stephan Nussberger, Werner Kühlbrandt

## Abstract

The TOM complex is the main entry point for precursor proteins into mitochondria. Precursor proteins containing targeting sequences are recognized by the TOM complex and imported into the mitochondria. We have determined the structure of the TOM core complex from *Neurospora crassa* by single-particle cryoEM at 3.3 Å resolution, showing its interaction with a bound presequence at 4 Å resolution, and of the TOM holo complex including the Tom20 receptor at 6-7 Å resolution. TOM is a transmembrane complex consisting of two β-barrels, three receptor subunits and three short transmembrane subunits. Tom20 has a transmembrane helix and a receptor domain on the cytoplasmic side. We propose that Tom20 acts as a dynamic gatekeeper, guiding precursor proteins into the pores of the TOM complex. We analyze the interactions of Tom20 with other TOM subunits, present insights into the structure of the TOM holo complex, and suggest a translocation mechanism.

## Introduction

The Translocase of the Outer Membrane (TOM) complex is an essential component of the outer membrane of mitochondria that acts as the main gate for protein import (1). As a result of the endosymbiotic origins of mitochondria, most of their proteins are produced in the cytosol as soluble precursors which are imported into the organelle (2, 3). Most of these proteins contain an N-terminal presequence that forms an amphipathic α-helix and acts as a mitochondrial targeting signal (MTS) (4, 5). Together with a diverse system of other import machineries, such as SAM in the outer membrane, TIM22 and TIM23 in the inner membrane, and multiple chaperones in the intermembrane space (IMS) and matrix, the TOM complex ensures that these proteins reach their final destination within the mitochondrion (6).

The TOM core complex of *Neurospora crassa* (NcTOM) in the mitochondrial outer membrane is a dimer with 5 subunits per monomer: Tom40, Tom22 and the small Toms (sT) Tom5, Tom6 and Tom7 (7). It has a mass of 148 kDa (8) and dimensions of roughly 130 Å by 100 Å (9). The protein translocation pores are formed by two copies of Tom40, each a 19-strand β-barrel with helical termini. Between the pores, two copies of the α-helical Tom22 span the membrane, with its disordered N and C-terminal domains facing the cytosol and IMS. The pronounced negative charge of the Tom22 N-terminus suggests that it interacts with the positive MTS of precursor proteins (10, 11). The small α-helical subunits Tom5, Tom6 and Tom7 are involved in complex assembly and are thought to play a role in complex stability and presequence recognition (12, 13).

In addition to the core complex components, the larger TOM holo complex contains the two receptor subunits Tom20 and Tom70 (14). Tom20 has a soluble core domain with 5 α-helices, including a tetratricopeptide repeat (TPR) at its C-terminus and a transmembrane helix at its N-terminus (15). It interacts hydrophobically with matrix-targeted precursor proteins and has been suggested to cooperate with Tom22 in the early steps of translocation (15, 16). Tom70 consists of 26 α-helices in a large soluble domain forming 11 TPR motives that are connected by a disordered loop to a transmembrane helix (17, 18). Tom70 specializes in carrier proteins and interacts with protein chaperones such as Hsp70 and Hsp90, but also cooperates with the mitochondrial import protein 1 (Mim1) in the biogenesis of membrane proteins (19, 20).

The structure of NcTOM has been investigated in negative-stain (7) and by electron cryo-microscopy (cryoEM), which yielded a 6.8 Å map (9). CryoEM structures of the yeast and human complex have been published at around 20 Å (21) and, more recently, at 3-4 Å resolution (22–24). The TOM holo complex is a challenging target because Tom20 and Tom70 are only loosely attached to the core complex (7, 25). Assembly, stoichiometry and interaction of Tom20 and Tom70 with the core subunits remain largely unknown. Recently, the structure of a TOM dimer with the Tom20 core domain chemically crosslinked to Tom40 has been reported, suggesting the presence of two copies of Tom20 per TOM dimer (26). We now set out to determine the high-resolution structure of the NcTOM core complex with bound pre-protein by single-particle cryoEM and to analyze the subunit composition of TOM holo complex through laser-induced liquid bead ion-desorption mass spectrometry (LILBID-MS). LILBID-MS is a native mass spectrometry technique that can examine intact as well as partially dissociated protein-complexes to identify subunit interactions (27, 28).

We present a 3.3 Å structure of the NcTOM core and a structure of NcTOM interacting with the positive MTS of rat aldehyde dehydrogenase visible at a lower density threshold. Furthermore, we present two structures showing the TOM core complex interacting with the peripheral components of Tom20 at about 7 Å resolution. Our map indicates that Tom20 is flexible, taking on two distinct conformations, with Tom22 as a docking platform.

## Results

### Isolation of the TOM holo complex

To investigate the structure of the TOM holo complex, we isolated mitochondria from *Neurospora crassa* hyphae with recombinant Tom22 containing a hexahistidine tag (9). We solubilized outer membrane vesicles (OMVs) in glyco-diosgenin (GDN) and isolated the complex by affinity purification and size-exclusion chromatography. Gel electrophoresis indicated that peak fractions (Fig. S1) contained all subunits of the TOM holo complex, including Tom20 and Tom70 (14). An additional band at around 30 kDa suggested the presence of the outer-membrane porin VDAC (29), a common contaminant in TOM preparations.

### Native Mass Spectrometry of TOM

We investigated the composition of the TOM holo complex and subunit interactions by LILBID-MS. Fig. 1 shows spectra up to 125,000 m/z, indicating the different subunits and fragment subcomplexes of the holo complex at two different laser intensities. Monomeric forms of the core subunits appear at high laser intensities (8), as seen in Fig. 1A. In addition, we identified the peaks for singly and doubly-charged Tom20 at 20,100 m/z and 10,200 m/z respectively. The predicted molecular mass of Tom20 is 20.23 kDa. The peak at 29,800 m/z can be assigned to the VDAC contaminant.

**Figure 1.**
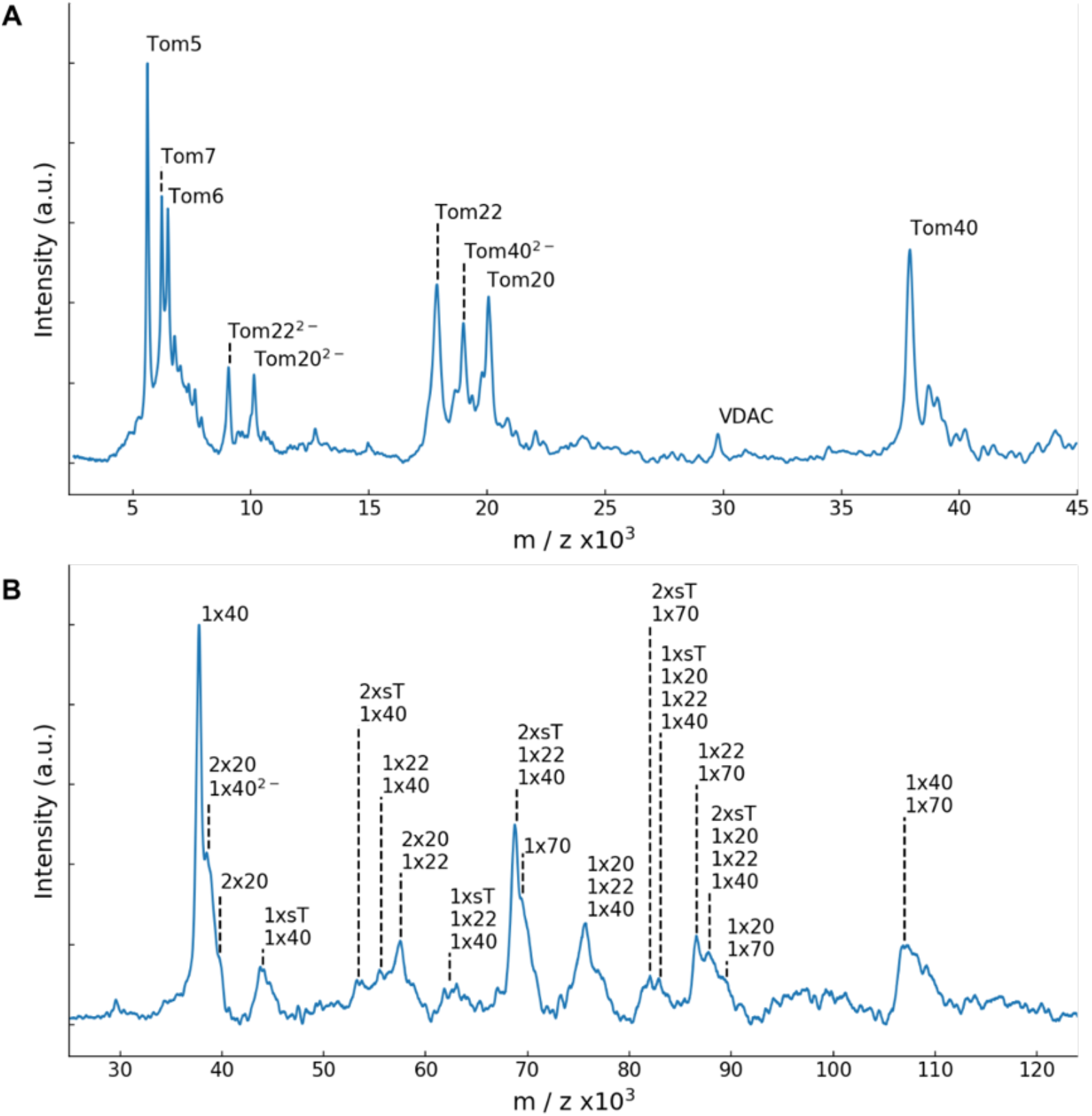
LILBID mass spectrometry of the TOM holo complex. Spectra of the complex at high (**A**) and low (**B**) laser intensities. Peaks were assigned according to their predicted molecular weights (ExPASy). (**A**) In the lower mass region, individual dissociated subunits are visible under harsh laser conditions. Tom5, Tom6, Tom7 appear as singly-charged entities, while Tom20, Tom22 and Tom40 carry one or two negative charges. A smaller peak was assigned to VDAC. (**B**) At reduced laser intensity, stable subcomplexes are visible in the mass range up to 125,000 m/z. Tom70 appears as a single molecule and forms subcomplexes with the small TOM subunits (sT), Tom20, Tom22 and Tom40. Tom20 forms subcomplexes with itself, with Tom22 and Tom40.

At reduced laser intensity (Fig. 1B) we observed peaks indicating larger assemblies. We identified the individual subunit Tom70 at 69,300 m/z, closely matching its predicted mass of 69.34 kDa. Additional peaks were assigned to subcomplexes formed by core subunits and holo receptors, revealing, for example, a stable interaction of Tom70 with two small Toms, labeled as sT_2_Tom70_1_. Likewise, Tom70 interacts with the translocation pore forming a Tom40_1_Tom70_1_ subcomplex, and with the receptor subunits forming the subcomplexes Tom20_1_Tom70_1_ and Tom22_1_Tom70_1_. We see evidence of a complex with Tom20_1_Tom22_1_Tom40_1_ stoichiometry, which contains also one or two small Toms, which might indicate a monomeric TOM holo assembly. This relates to the subcomplexes formed by Tom22, Tom40 and a variable number of sT, as reported (8). Interestingly, we see Tom20_2_ dimers, also forming subcomplexes with other subunits, suggesting the presence of two copies of Tom20 per complex. Previously, the crystal structure of Tom20 with bound presequence (PDB 2V1S) was reported to be a dimer, although dimer formation was reported as biologically irrelevant, due to the nature of the residues involved in the interaction (30). Peak assignment based on the subunits mass can be found in table S1.

### Structure of the dimeric TOM core complex

To study the TOM translocation mechanism, we incubated the purified holo-complex with a synthetic peptide containing the presequence of rat aldehyde dehydrogenase (pALDH). We then plunge-froze the mixture and performed single-particle analysis of the complex in GDN, which resulted in a 3.32 Å resolution map of the TOM core complex to which we applied C2 symmetry during the final reconstruction step (Fig. 2A and Fig. S2). We identified the 5 core subunits of NcTOM based on our published 6.8 Å structure of the dimeric core complex (9). Our high-resolution TOM core map is composed of two copies of the Tom40 barrel, the helical Tom22, Tom5, Tom6 and Tom7 subunits, but it lacks the receptors Tom20 and Tom70, and the presequence.

**Figure 2.**
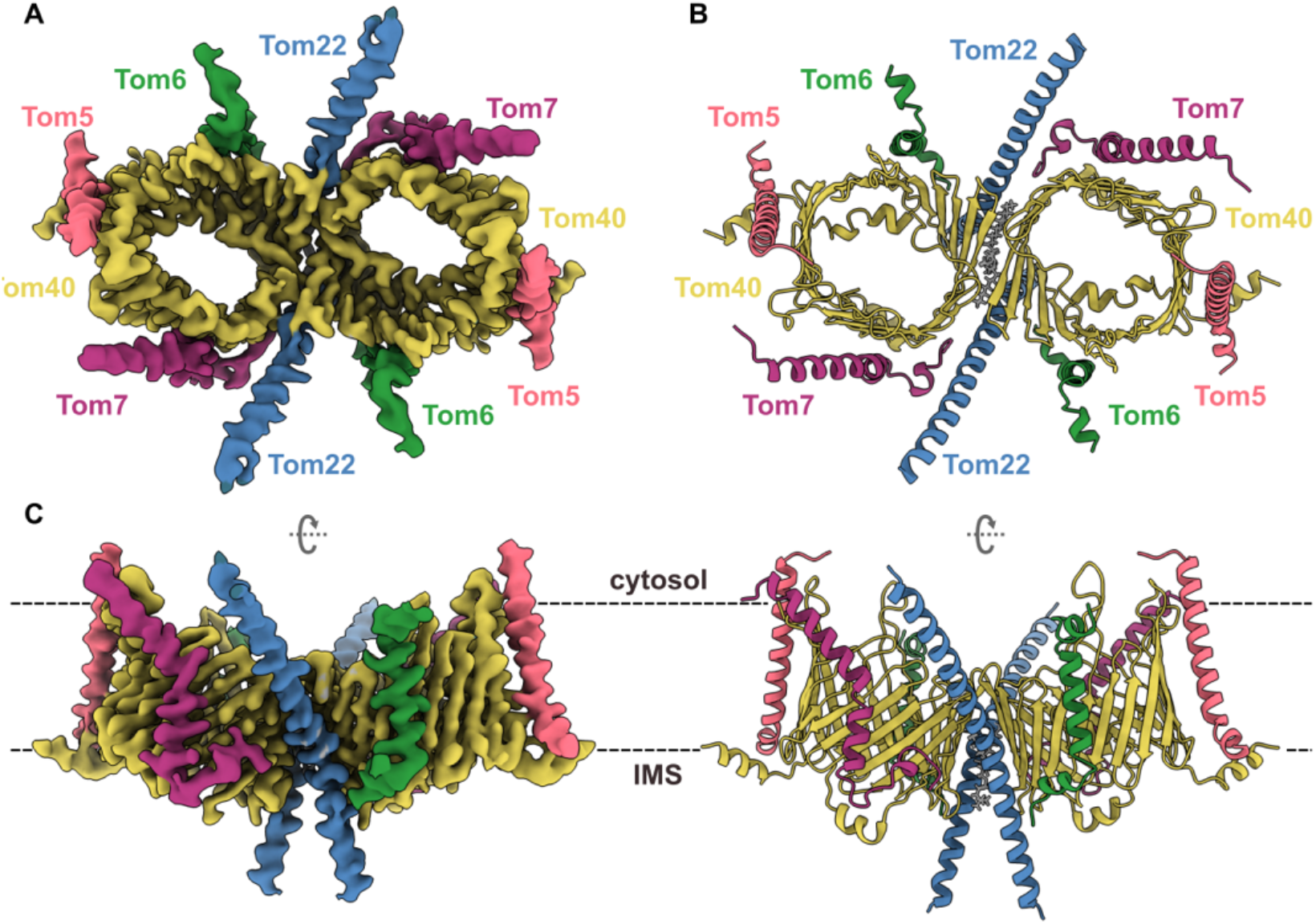
CryoEM map and model of the TOM core complex from *Neurospora crassa*. Subunit assignment of the components of the NcTOM core complex. Tom40, yellow; Tom22, blue; Tom5, Tom6 and Tom7 are pink, green and purple, respectively. (**A**) TOM complex dimer at 3.32 Å resolution seen from the cytosol. (**B**) Cartoon representation of the atomic model, including a lipid molecule between the two pores seen from the cytosol. (**C**) Side view of the map and model from the outer mitochondrial membrane. The approximate position of the lipid bilayer is indicated by dashed lines.

We built an atomic model (Fig 2B and Fig. S3), starting from a prediction model generated with AlphaFold-Multimer. Our map contains clear densities for the 19 individual β-strands of the Tom40 barrel and the loops connecting them (Fig. S4). The longest loop, joining strands 14 and 15, is visible at a lower density threshold, indicating that it is flexible. As observed in our previous model of Tom40 (PDB 5O8O), the N-terminus starts with two short helices (a1 and a2) and the C-terminus ends in one helix (a3). Helix a1 starts outside the pore and interacts with Tom5 at the IMS. After an unstructured but highly conserved stretch, it turns into helix a2 inside the Tom40 pore, interacting with β-strands 11 to 16. Internal short helices such as a2 are common features of membrane-embedded β-barrels, and their mutation or deletion can lead to structural reshaping and destabilizing of the barrels (31, 32). At the end of β-strand 19, a3 extends into the IMS and folds back into the pore to interact with strand 4. Due to flexibility, a3 is only visible at a lower density threshold, while its C-terminal residues are well-ordered. At the point of contact between the monomers, we see interactions between strands 19, 1 and 2 of the two Tom40 barrels.

As we found previously, the two TOM core monomers sit in the membrane at an angle of ~20°, so that the pores are tilted relative to each other (9). Between them, we identified a phospholipid that interacts with Tom40 and Tom22 (Fig. 3A). The lipid is in contact with strands 17, 18 and 19 of both Tom40s, and is likely stabilized by residue F309, for which we see two rotamer conformations (Fig. 3B). The map shows another four elongated, bulky densities on each side of the dimer, in close contact with Tom40 and Tom22. We assigned these densities that span half the membrane to GDN molecules, the detergent used for solubilization (Fig. 3C). Nearly 70% of the mitochondrial outer membrane is accounted for by phosphatidylcholine and phosphatidylethanolamine (33); we propose that the remaining unmodeled elongated densities around the dimer in the map correspond to these lipids.

**Figure 3.**
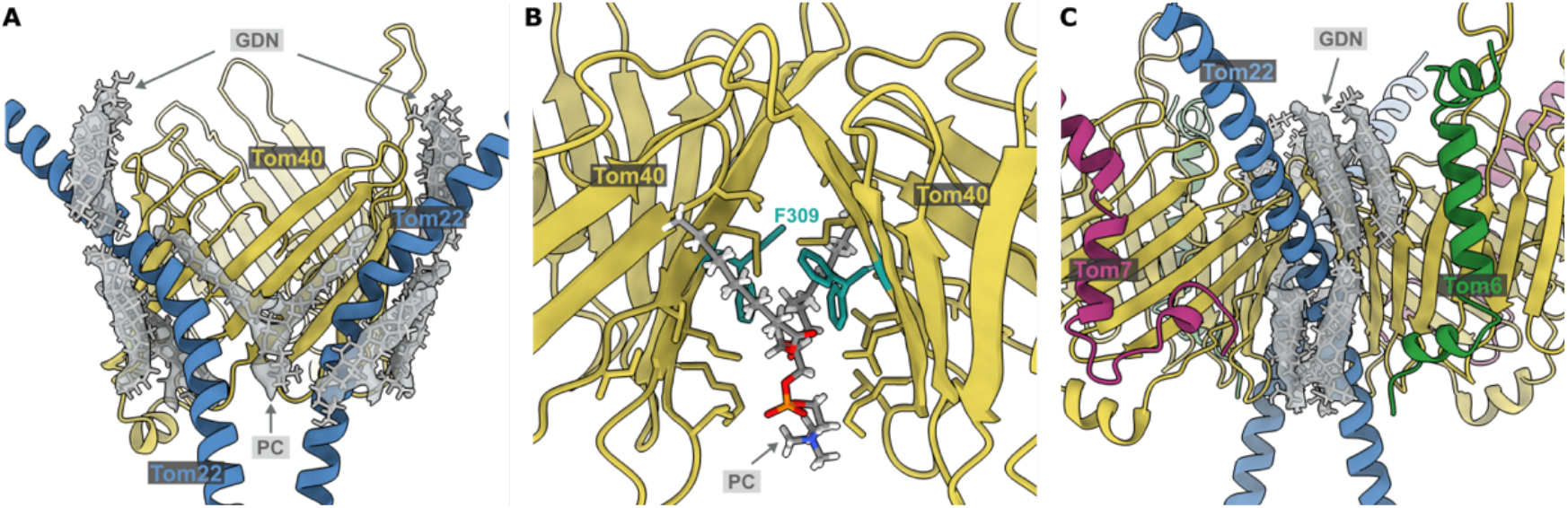
Lipid densities surrounding Tom22 and Tom40. (**A**) Two molecules interact with the TOM core complex: phosphatidylcholine (PC) and glycol-diosgenin (GDN). (**B**) Close-up of PC at the dimer interface, where we see two rotamer conformations for F309 in Tom40. (**C**) Four molecules of GDN were identified around each monomer of the complex.

Our model also contains two copies of the Tom22 transmembrane helix. This subunit, which extends from the cytosol towards the IMS, is only partially resolved, as its hydrophilic N- and C-termini are disordered (Fig. 2C). They were shown to interact with presequences and with other subunits (34). Similarly, the flexible N-termini of Tom5 and Tom6 are not visible, while their transmembrane domains are clearly helical. In our map, Tom7 is mostly complete, embedded in the membrane and displaying its characteristic Z shape. Our assignment of NcTOM subunits matches our earlier 6.8 Å map (EMD-3761), except for the IMS domain of Tom6, which appeared to be longer in our previous map (Fig. S5), perhaps due to anisotropic resolution.

### Precursor protein translocation

Refinement of the same single-particle dataset with limited alignment resolution yielded a C2-symmetrical 4 Å map, in which each of the translocation pores contained a rod-shaped density crossing the Tom40 barrel (Fig. 4), visible at lower thresholds. A difference map (Fig. S6) against a map generated from our NcTOM core model indicated that the rod-shaped densities are consistent with the dimensions of an α-helix. We propose they correspond to the precursor substrate pALDH that was incubated with our complex, which thus appears to be captured in translocation. We rigid-body-fitted an AlphaFold prediction of our pALDH construct into the density, using UCSF ChimeraX.

**Figure 4.**
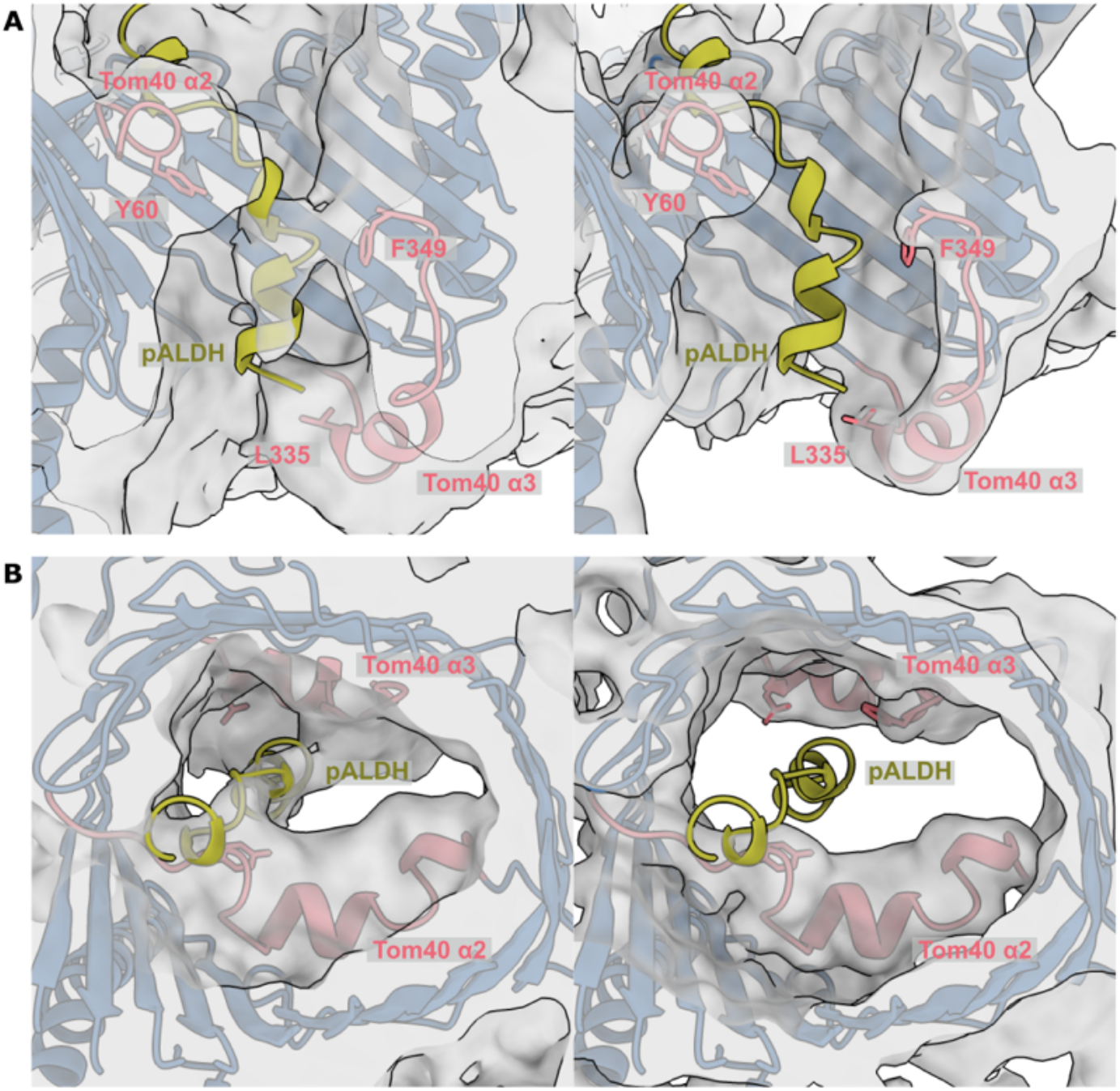
Presequence bound to Tom40. Transversal cuts of the model-map superposition showing the presequence density inside the translocation pore at lower (left) and higher (right) density thresholds. The pALDH structure generated with AlphaFold is shown in yellow, the 4 Å map of the TOM core complex in gray, and the two inner helices of Tom40 in pink. (**A**) Side view of the inner Tom40 pore showing Y60, F349 and L335. The presequence density spans from the cytoplasmic side towards the IMS, interacting on its way with α2 and α3. (**B**) View of the pore from the cytosol.

The map in Fig. 4 shows the interaction of the helical presequence, superimposed on our TOM core C2 model. Extending from the cytosol into the IMS, the presequence density makes contact with α2 and α3 inside Tom40, interacting with regions rich in hydrophobic residues. The precursor substrate helix interacts with α2 near Y60 of Tom40. On the opposite side it approaches F349, the C-terminal residue of Tom40 at the end of α3. The presequence density further appears to interact with α3 at the IMS side of TOM within close range of L335. This suggests a possible involvement of Y60 and L335 in presequence recognition and translocation.

### Interaction of Tom20 with the TOM core complex

We extracted more information on the cytosolic domains of Tom20 and Tom22 from the same dataset at lower resolution. After two rounds of 3D classification and refinement, we resolved two maps that we assign to two different conformations of Tom20, C_1_ and C_2_, at 6.7 Å and 6.6 Å resolution, respectively (Fig. S2 and S7). The rod-shaped density of an α-helix protrudes from the edge of the micelle and appears to be suspended over the pores. At its end, a globular domain becomes visible at a lower density threshold.

We were able to rigid-body-fit the AlphaFold prediction models for Tom20 and Tom22 into both maps (Fig. S8). As shown in Fig. 5, we observe that one Tom22 helix bends further to the side of the core complex on the cytosolic side, as in the *H. sapiens* core complex (24), and stretches across the asymmetric micelle. The transmembrane helix of Tom20 emerges from the micelle and interacts with Tom22 at the membrane surface. A helix, with its connected receptor domain, appears to hover above of the translocation pores, roughly parallel to the membrane plane. At the interface, part of the N-terminus of Tom22 wraps around Tom20, connecting it to the docking site.

**Figure 5.**
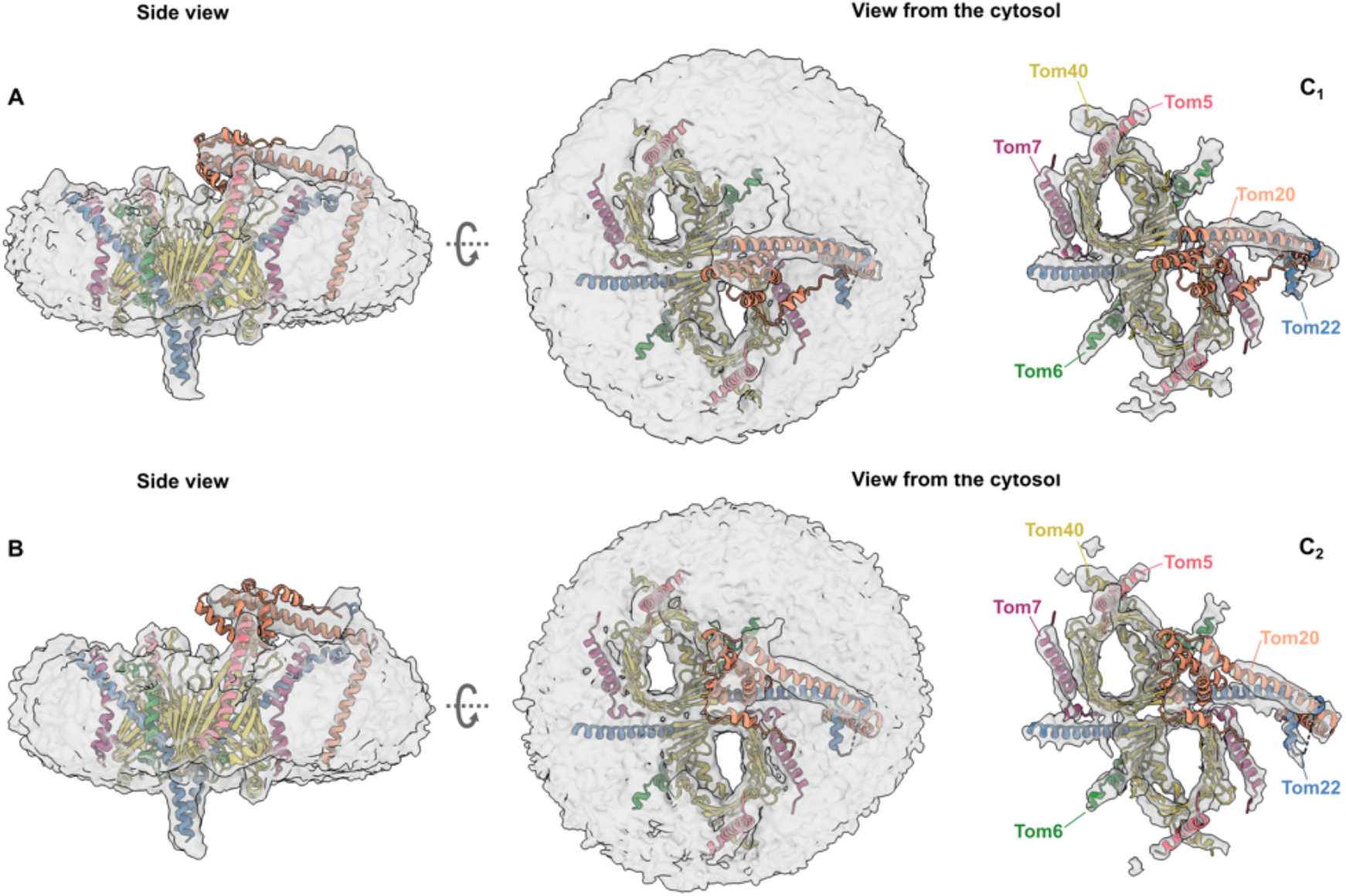
Tom20 assumes two discrete conformations on the cytosolic side. Fit of an AlphaFold model of Tom20 to the cryoEM map of the TOM holo complex at 6.7 Å (**A**) and 6.6 Å (**B**), superimposed with the core model. The cytosolic domain of Tom20 (orange) assumes two distinct positions (C_1_ and C_2_) aligned with Tom22 at the cytosol-membrane in the center of the TOM core dimer or closer to one pore, where it appears to interact with Tom6.

The two resolved maps differ on the position of Tom20 on the cytoplasmic side of the complex. In Fig. 5A, Tom20 is situated right on top and in between the translocation pores (C_1_), while in Fig. 5B, Tom20 leans towards one of the pores within close range of Tom6 and the loop connecting Tom40 strands 14 and 15 (C_2_). This might indicate that the receptor domain of Tom20 serves both pores and can shuttle between them.

## Discussion

We used cryoEM and LILBID-MS to gain insight into the structure and translocation mechanism of the TOM complex. Our TOM core map shows a conserved structure in close agreement to the previously published cryoEM structures of *N. crassa*, yeast and human TOM. However, specific differences between species are evident (Fig. S9). The C-terminus of NcTom40 is a flexible helix that extends into the IMS and folds back into the translocation pore, while in the human complex, this helix is replaced by a longer Tom7 C-terminus. In yeast, a corresponding helix is instead oriented towards Tom22 (35). These differences are visible in the IMS exit pathway and would affect the interaction of the complex with presequences. Similarly, the Tom40 loop between strands 14 and 15 is considerably longer in *N. crassa* than in the human complex, which might influence presequence insertion into the pore. We confirm the presence of a phospholipid at the interface between the two copies of Tom40, which might serve to maintain the tilt angle between the two Tom40 barrels (22). The four elongated strong densities around each Tom22 that we assign to GDN molecules are bound to be occupied by lipids in the membrane.

Our lower-resolution presequence density inside the pores is consistent with earlier research, suggesting that Tom40 uses a combination of acidic and hydrophobic patches to translocate the presequence towards the IMS (36). Crosslinking studies have indicated that other presequences bind to the cytoplasmic side of Tom40 (37), however they most likely relate to early steps in translocation. Our density appears in the center of the pore close to the IMS exit site, and may thus reflect a late stage of the translocation process (Fig. S6). This interaction points to the trans-binding site within Tom40, supported by Tom7 and Tom22 (38, 39). The presequence is likely to adopt multiple positions in the complex. This would explain why it appears at a low density threshold in the cryoEM map, which is an average of all TOM core particles with bound presequence.

Our results shed new light on the function of the receptor Tom20. In contrast to an earlier study with the human TOM complex (26), we neither crosslinked Tom20 to the core complex nor did we impose twofold symmetry. Our map is therefore likely to represent the native structure, which is clearly asymmetric and thus appears to have only one copy of Tom20 in the holo-complex. Our LILBID-MS results show multiple subcomplexes composed of Tom20, Tom22 and Tom40 with different stoichiometries (Fig. 1B). This is consistent with the cytosolic domain of Tom20 being flexible, capable of taking on different conformations in its interactions with the TOM core complex. Our maps indicate a strong interaction of the acidic patch of the N-terminus of Tom22 with a positively charged patch in the Tom20 helix (Fig. S10), confirming that Tom22 is a docking point required for optimizing the receptor function of Tom20 (16, 34, 40). Based on our models, we propose that Tom20 docks to the Tom22 helix at the cytosolic membrane surface, while Tom22 and Tom40 bind strongly to each other through sidechain-specific hydrophobic and electrostatic contacts (Fig. 6).

**Figure 6.**
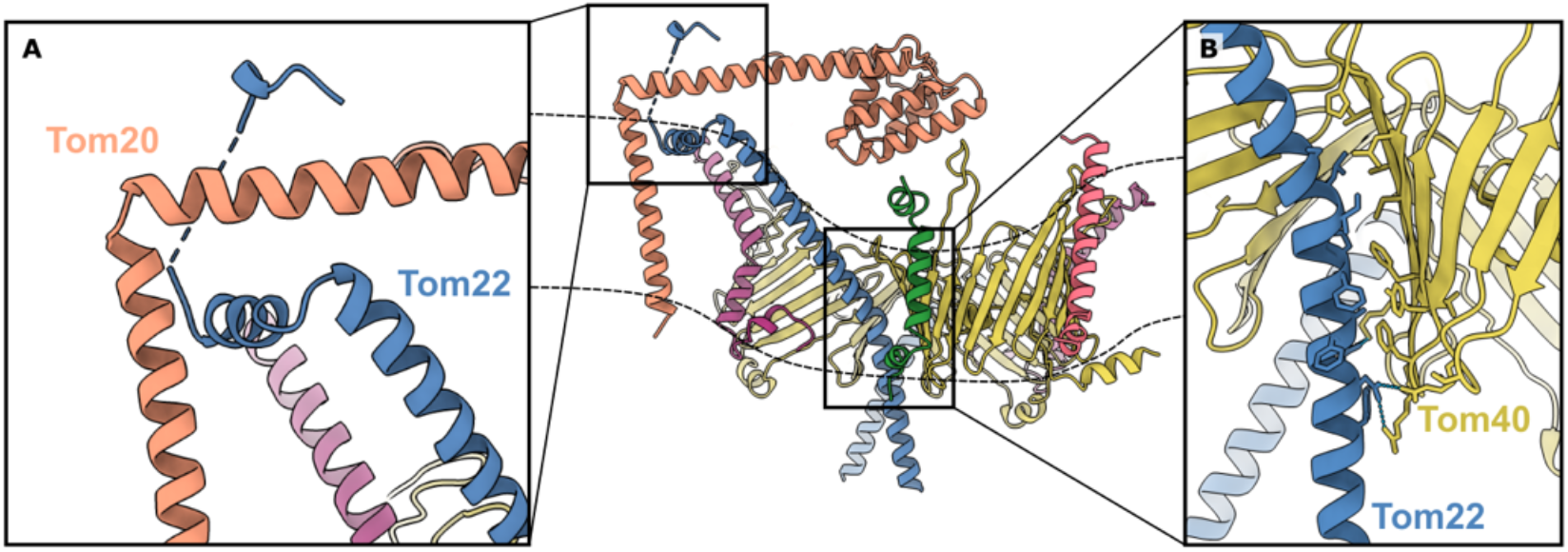
Interactions between Tom20, Tom22 and Tom40. Proposed interactions between Tom20 in orange, Tom22 in blue and Tom40 in yellow, based on our core and holo models. The position of the outer mitochondrial membrane is indicated by dashed lines. (**A**) Rigid-body-fitted model of Tom20 docked to Tom22. Tom20 is stabilized by the N-terminus of Tom22 wrapping around it. (**B**) Tom22 and Tom40 are held together in the membrane by hydrophobic and electrostatic contacts.

We propose that in the native holo complex Tom20 takes on two alternative conformations. In conformation C_1_, Tom20 lies parallel to the membrane, roughly aligned to Tom22, within close range of both translocation pores (Fig. 7A). In conformation C_2_, the Tom20 receptor domain approaches one of the pores and interacts with Tom40 and perhaps Tom6 (Fig. 7B,). This second conformation agrees with crosslinking studies that indicate interaction of the longest Tom40 loop between strands 14 and 15 with Tom20 (37), and the LILBID-MS subcomplexes formed by Tom20, Tom22, Tom40 and at least one small TOM subunit (Fig. 1B). Moreover, conformation C_2_ is consistent with the model predicted by AlphaFold for the Tom20_1_Tom22_1_Tom40_1_ subcomplex (Fig. S8A). Additionally, other crosslinking studies have indicated the interaction of a presequence with Tom6 in the early stages of translocation (41). Taking all this into consideration, we suggest that, upon contact with the presequence, Tom20 approaches Tom6, interacts with Tom40, and deposits the presequence into the pore, initiating the translocation process.

**Figure 7.**
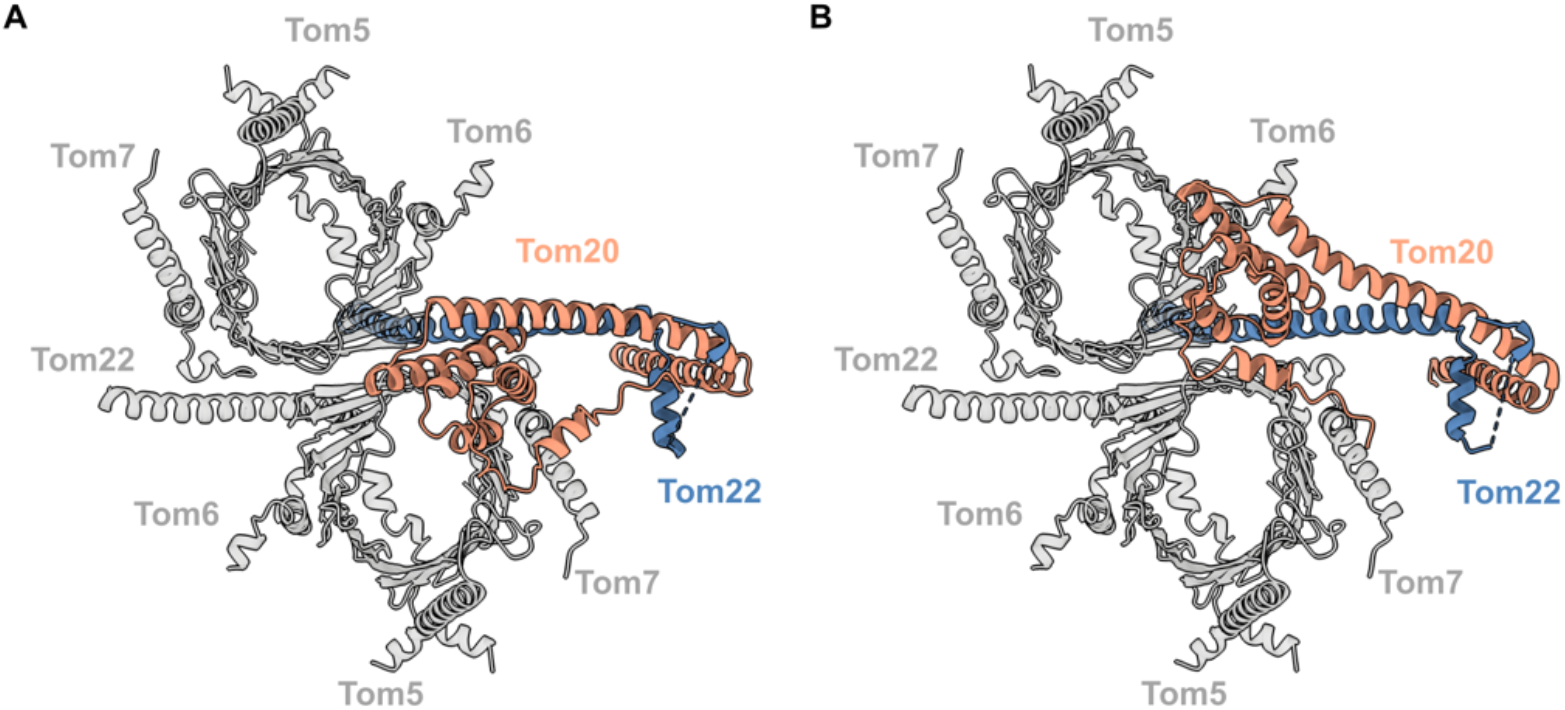
Model of the TOM complex including Tom20. Cartoon representation of Tom20 (orange), docked on Tom22 (blue), interacting with the TOM core complex (gray), as seen from the cytosol in two conformations. In conformation C_1_ (**A**), Tom20 assumes a central position between the two pores, while in C_2_ (**B**), Tom20 approaches one pore and interacts with Tom40 near Tom6.

Our non-symmetrized map indicates one copy of Tom20 in the complex, although our LILBID-MS spectra suggest that a fraction of TOM subcomplexes can contain two copies of Tom20 (Figure 1B). However, a minor population of TOM complexes may not be detected as a separate class in image processing. We nevertheless attempted to fit two copies of Tom20 in its two different conformations into our TOM core model, but found that the two receptor domains would come into close contact or clash on the cytoplasmic side. A steric clash would make it unlikely that two copies of Tom20s can coexist simultaneously in one TOM dimer (Fig. S11).

We see a peak corresponding to Tom70 in our LILBID-MS spectra (Figure 1B), as well as subcomplexes formed by Tom70 in association with Tom20, Tom22, Tom40 and up to two small subunits, but we are not able to assign it to any particular density region. The interaction between Tom20 and Tom70 has been reported before in vitro in other organisms by crosslinking (42). However, a stable interaction might depend on Tom70 binding a precursor protein. More work is required to establish the position of Tom70 relative to the core complex and to fully understand their function.

In conclusion, we propose a translocation pathway for the TOM core complex that includes a flexible Tom20 in its two observed conformations (Movie S1). We propose that Tom22 not only acts as a presequence receptor, but serves as a docking platform for Tom20, stabilizing Tom20 through electrostatic interactions at its N-terminus. Apparently Tom20 can reach both pores from conformation C_1_ (Fig. 7A). We propose that, upon contact with a presequence, Tom20 changes its conformation to C_2_ (Fig. 7B), hovering above one pore where it interacts with the Tom40 loop between strands 14 and 15. In this way, Tom20 would deliver the presequence to Tom40, which in turn translocates the precursor protein by means of acidic and hydrophobic interactions.

## Materials and Methods

### Growth of Neurospora crassa and preparation of mitochondrial outer membrane vesicles

*Neurospora crassa* (strain GR-107) containing a hexahistidinyl-tagged form of Tom22 was cultured and mitochondria were isolated as described (43). Briefly, ~1.5 kg wet weight of hyphae was homogenized in 250 mM sucrose, 2 mM EDTA, 20 mM Tris pH 8.5, 1 mM phenyl methylsulfonyl fluoride (PMSF) in a Waring mixer at 4 °C. ~1.5 kg of silica sand was added and cell walls were broken by passing the suspension through a corundum mill. Cellular residues were pelleted and discarded in two centrifugation steps (4000 × g) for 5 min at 4 °C. Mitochondria were sedimented in 250 mM sucrose, 2 mM EDTA, 20 mM Tris pH 8.5, 1 mM PMSF at 17,000 × g for 80 min. This step was repeated to improve purity. The isolated mitochondria were suspended in 250 mM sucrose, 20 mM Tris pH 8.5, 1 mM PMSF with a final protein concentration of 50 mg/ml. Mitochondrial membranes were then separated from soluble matrix and intermembrane space proteins by centrifugation at 17,700 × g after mitochondria were swollen at a protein concentration of 2 mg/ml in 5 mM Tris pH 8.5, 1 mM EDTA, and 1 mM PMSF. To obtain the outer membranes of the mitochondria, the membrane pellets were resuspended in the same buffer and homogenized in an automated glass-Teflon douncer for 60 min at 4°C to separate the outer from the inner membranes.

Outer membranes (OMVs) were isolated by sedimentation and flotation centrifugation of the homogenate in sucrose step gradients in 20 mM Tris pH 8.5, 1 mM EDTA and 1 mM PMSF as previously described (14). The isolated outer membranes were diluted threefold with 50 mM Tris pH 8.5, sedimented by centrifugation at 250,000 × g, resuspended in 50 mM Tris pH 8.5 with a protein concentration of ~1 mg/ml and used directly to isolate the TOM holo complex.

### Purification of the TOM holo complex

The TOM holo complex was isolated from OMVs in solubilization buffer (20% glycerol, 10mM MOPS pH 7.0, 50 mM potassium acetate, 50 mM imidazole, 1 mM PMSF and 1% GDN) at a protein concentration of 1 mg/ml. After incubation for 1 h at 4°C, the lysate was centrifuged at 13,000 x g for 20 min. The clarified extract was loaded onto a nickel-nitrilotriacetic acid (Ni-NTA) column. Non-specifically bound proteins were washed off using 10mM MOPS pH 7.0, 50 mM potassium acetate, 50 mM imidazole, 1 mM PMSF and 0.02% GDN). The complex was eluted with 300 mM imidazole in the same buffer and concentrated (AmiconUltra 100 kDa cutoff). The purity of the eluted fractions was assessed by SDS-PAGE and Coomassie Brilliant Blue staining.

For LILBID-MS we further purified TOM holo complex using a Superdex 200 Increase size exclusion column (GE) equilibrated with 10 mM Tris pH 7.0, 15 mM ammonium acetate and 0.02% GDN. After SDS-PAGE, the peak fractions containing the TOM holo complex were pooled together and concentrated to 4 mg/ml (AmiconUltra 100 kDa cutoff). For CryoEM, the complex was incubated for 1 h with excess pALDH at a 1:8 ratio (44). The mix was loaded onto a Superdex 200 Increase size exclusion column (GE) equilibrated with 50 mM KPO_4_ pH 8.0, 50 mM KCl, 1 mM EDTA, 1 mM TCEP and 0.02% GDN. Fractions were assessed by SDS-PAGE and Coomassie Brilliant Blue staining.

### Laser Induced Liquid Bead Ion Desorption Mass Spectrometry

For LILBID-MS analysis, ions were generated with an IR laser from 50 μm microdroplets containing the proteins of interest (45). Microdroplets were produced by a commercially available piezo-driven droplet generator (MD-K-130; Microdrop Technologies GmbH). The IR laser operated at the absorption wavelength of water (2.94 μm) and droplets were produced and irradiated at a frequency of 10 Hz. The IR-laser power was varied in a range of 10 mJ - 23 mJ. Ions were detected with a home-built time-of-flight analyzer, operating at a vacuum of 10-6 mbar. Each measurement was performed in negative ion mode with a sample volume of 4 μl. All shown mass spectra were normalized to 1 and represent averaged signals of 1000 droplets. Spectra were analyzed and data were processed with Massign (46). Peaks were assigned on the basis of predicted average molecular mass of the individual subunits, calculated using ExPASy (47) (see table S1 for details).

### Precursor protein preparation

The precursor peptide pALDH was synthesized by GenScript, resuspended in H2O upon delivery, aliquoted and frozen until used. The peptide consisted of the first 19 amino acids of the MTS of rat aldehyde dehydrogenase plus a StrepII-tag joined by a linker to its C-terminus: MLRAALSTARRGPRLSRLLSGGGSWSHPQFEK.

### CryoEM specimen preparation and data acquisition

A TOM holo complex peak fraction containing ~ 2 mg/ml protein was used for cryoEM. Roughly 3 μl were applied to a glow-discharged C-Flat 1.2/1.3 Cu grid, blotted for 3 s at 100% humidity at 4°C and flash-frozen in a Vitrobot Mark IV (FEI). Images were recorded at 300 kV using a Titan Krios electron microscope (ThermoFisher Scientific) equipped with a Gatan K3 camera in counting mode and a Gatan BioQuantum energy filter. Dose-fractionated movies were acquired with 3 s exposure at a 105,000x nominal magnification, resulting in a pixel size of 0.83 Å. The total accumulated dose was 55 e^-^/A^2^. Image defocus was in the range of −1.2 to - 3.0 μm.

### Single-particle analysis of the TOM core complex

Images were processed using Relion-4.0 (48). Movies were motion-corrected using MotionCor2 (49) and CTF parameters were initially estimated using CTFFIND-4 (50), both as implemented in Relion. A particle-picking model was manually built using crYOLO (51) and subsequently applied to the entire dataset. After extraction, 2D classification was used to discard artefacts, and a set of the 620,000 best particle images was separated by 3D classification. Initial 3D refinement indicated a resolution of 4.2 Å with Bayesian polishing. The polished particles were imported into cryoSPARC v3 (52), where nonuniform refinement produced a map at 3.6 Å resolution. After a round of 3D classification without alignment, 304,000 particles were subjected to non-uniform refinement with imposed C2 symmetry to 3.37 Å resolution. Local refinement in CryoSPARC with a mask around the entire core complex improved the resolution to 3.32 Å, as assessed by the gold-standard FSC = 0.143 criterion. Local resolution was determined by cryoSPARC (see Fig. S2 and table S2 for details). The same particles were non-uniformly refined with a 10 Å maximum align resolution limit and imposed C2 symmetry in cryoSPARC. The refined map had a resolution of 4 Å according to the gold-standard FSC, and was used to study the preprotein bound TOM core complex. The presequence density was identified by subtraction of a map generated from the TOM core model using UCSF ChimeraX (53) (Fig. S6). The detergent micelle was deleted from the difference map using the volume eraser.

### Single-particle analysis of the TOM core + Tom20 complex

Polished particles were further processed in Relion. A mask covering the area assigned to Tom20 was created in UCSF ChimeraX. Following masked 3D classification without alignment, two classes with distinct Tom20 conformations were selected, with 140,000 and 120,000 particles each, and independently refined without enforced symmetry to resolutions of 6.6 and 6.7 Å respectively (see Fig. S2 and S7 for details).

### Model building

Atomic model building of the TOM core complex was based on the AlphaFold-Multimer (54) prediction of the core dimer, then fitted into the refined map using Coot (55) and ISOLDE (56) within UCSF ChimeraX. Additional real-space refinement was performed in Phenix (57). Phosphatidylcholine and diosgenin were and fitted into the structure. The structure of the pALDH construct and the oligomer formed by Tom20, Tom22 and Tom40 were predicted using AlphaFold-Multimer. The preprotein bound TOM core complex was rigid-body-fitted with UCSF ChimeraX. The TOM core plus Tom20 model was based on the AlphaFold (58) predictions of Tom20 and Tom22 (Fig. S8). For each conformation, Tom20 and Tom22 were rigid-body-fitted to the map, relaxed using Coot and then merged into the TOM core model (see table S2 for more information).

## Acknowledgments

We thank Beate Nitschke for help with mitochondria preparation and Parinaz Lordifard for her contribution in sample preparation. We thank Janet Vonck and Noor Agip for advice on modelling. We thank Juan Castillo and Özkan Yildiz for computational support, and Max Bernhard for critical comments on the manuscript.

## Author Contributions

PO and TB designed the experiments. SN and PO purified and characterized the TOM complex. JM and NM carried out the mass spectrometry analysis. PO built the models, analyzed the results and wrote the original draft. WK supervised the project. All authors reviewed and edited the paper.

## Data availability

The cryoEM map and atomic model of the TOM core complex have been deposited in the Protein Data Bank and in the Electron Microscopy Data Bank with accession codes PDB 8B4I and EMD-15849. The cryoEM maps of the two conformations of the TOM complex with Tom20 have been deposited in the Electron Microscopy Data Bank with accession codes EMD-15850 and EMD-15856. All data needed to evaluate the conclusions are present in the paper and/or the Supplementary Materials.

## References

1. O. Schmidt, N. Pfanner, C. Meisinger, Mitochondrial protein import: from proteomics to functional mechanisms. Nat Rev Mol Cell Biol 11, 655–667 (2010).

2. S. G. Garg, S. B. Gould, The Role of Charge in Protein Targeting Evolution. Trends in Cell Biology 26, 894–905 (2016).

3. M. C. Avendaño-Monsalve, J. C. Ponce-Rojas, S. Funes, From cytosol to mitochondria: the beginning of a protein journey. Biological Chemistry 401, 645–661 (2020).

4. D. Roise, S. J. Horvath, J. M. Tomich, J. H. Richards, G. Schatz, A chemically synthesized pre-sequence of an imported mitochondrial protein can form an amphiphilic helix and perturb natural and artificial phospholipid bilayers. The EMBO Journal 5, 1327–1334 (1986).

5. A. Chacinska, C. M. Koehler, D. Milenkovic, T. Lithgow, N. Pfanner, Importing Mitochondrial Proteins: Machineries and Mechanisms. Cell 138, 628–644 (2009).

6. N. Wiedemann, N. Pfanner, Mitochondrial Machineries for Protein Import and Assembly. Annu. Rev. Biochem. 86, 685–714 (2017).

7. U. Ahting, et al., The Tom Core Complex. Journal of Cell Biology 147, 959–968 (1999).

8. F. Mager, L. Sokolova, J. Lintzel, B. Brutschy, S. Nussberger, LILBID-mass spectrometry of the mitochondrial preprotein translocase TOM. J. Phys.: Condens. Matter 22, 454132 (2010).

9. T. Bausewein, et al., Cryo-EM Structure of the TOM Core Complex from Neurospora crassa. Cell 170, 693–700.e7 (2017).

10. M. Moczko, et al., The intermembrane space domain of mitochondrial Tom22 functions as a trans binding site for preproteins with N-terminal targeting sequences. Mol Cell Biol 17, 6574–6584 (1997).

11. J. Brix, S. Rüdiger, B. Bukau, J. Schneider-Mergener, N. Pfanner, Distribution of Binding Sequences for the Mitochondrial Import Receptors Tom20, Tom22, and Tom70 in a Presequence-carrying Preprotein and a Non-cleavable Preprotein. Journal of Biological Chemistry 274, 16522–16530 (1999).

12. T. Becker, et al., Assembly of the Mitochondrial Protein Import Channel. MBoC 21, 3106–3113 (2010).

13. K. Dietmeier, et al., Tom5 functionally links mitochondrial preprotein receptors to the general import pore. Nature 388, 195–200 (1997).

14. K.-P. Künkele, et al., The Preprotein Translocation Channel of the Outer Membrane of Mitochondria. Cell 93, 1009–1019 (1998).

15. Y. Abe, et al., Structural Basis of Presequence Recognition by the Mitochondrial Protein Import Receptor Tom20. Cell 100, 551–560 (2000).

16. K. Yamano, et al., Tom20 and Tom22 Share the Common Signal Recognition Pathway in Mitochondrial Protein Import. Journal of Biological Chemistry 283, 3799–3807 (2008).

17. D. E. Gordon, et al., Comparative host-coronavirus protein interaction networks reveal pan-viral disease mechanisms. Science 370, eabe9403 (2020).

18. Y. Wu, B. Sha, Crystal structure of yeast mitochondrial outer membrane translocon member Tom70p. Nat Struct Mol Biol 13, 589–593 (2006).

19. J. C. Young, N. J. Hoogenraad, F. U. Hartl, Molecular Chaperones Hsp90 and Hsp70 Deliver Preproteins to the Mitochondrial Import Receptor Tom70. Cell 112, 41–50 (2003).

20. T. Becker, et al., The mitochondrial import protein Mim1 promotes biogenesis of multispanning outer membrane proteins. Journal of Cell Biology 194, 387–395 (2011).

21. K. Model, C. Meisinger, W. Kühlbrandt, Cryo-Electron Microscopy Structure of a Yeast Mitochondrial Preprotein Translocase. Journal of Molecular Biology 383, 1049–1057 (2008).

22. Y. Araiso, et al., Structure of the mitochondrial import gate reveals distinct preprotein paths. Nature 575, 395–401 (2019).

23. K. Tucker, E. Park, Cryo-EM structure of the mitochondrial protein-import channel TOM complex at near-atomic resolution. Nat Struct Mol Biol 26, 1158–1166 (2019).

24. W. Wang, et al., Atomic structure of human TOM core complex. Cell Discov 6, 1–10 (2020).

25. P. J. T. Dekker, et al., Preprotein Translocase of the Outer Mitochondrial Membrane: Molecular Dissection and Assembly of the General Import Pore Complex. Mol Cell Biol 18, 6515–6524 (1998).

26. J. Su, et al., Structural basis of Tom20 and Tom22 cytosolic domains as the human TOM complex receptors. Proc. Natl. Acad. Sci. U.S.A. 119, e2200158119 (2022).

27. O. Peetz, et al., LILBID and nESI: Different Native Mass Spectrometry Techniques as Tools in Structural Biology. J. Am. Soc. Mass Spectrom. 30, 181–191 (2019).

28. J. Mezhyrova, et al., Membrane insertion mechanism and molecular assembly of the bacteriophage lysis toxin ΦX174-E. FEBS J 288, 3300–3316 (2021).

29. S. Schmitt, et al., Proteome analysis of mitochondrial outer membrane from Neurospora crassa. Proteomics 6, 72–80 (2006).

30. T. Saito, et al., Tom20 recognizes mitochondrial presequences through dynamic equilibrium among multiple bound states. The EMBO journal 26, 4777–4787 (2007).

31. H. Naveed, R. Jackups, J. Liang, Predicting weakly stable regions, oligomerization state, and protein–protein interfaces in transmembrane domains of outer membrane proteins. Proceedings of the National Academy of Sciences 106, 12735–12740 (2009).

32. R. Böhm, et al., The Structural Basis for Low Conductance in the Membrane Protein VDAC upon β-NADH Binding and Voltage Gating. Structure 28, 206–214.e4 (2020).

33. G. Hallermayer, W. Neupert, Lipid Composition of Mitochondrial Outer and Inner Membranes of Neurospora crassa. Biological Chemistry 355, 279–288 (1974).

34. S. van Wilpe, et al., Tom22 is a multifunctional organizer of the mitochondrial preprotein translocase. Nature 401, 485–489 (1999).

35. T. Bausewein, H. Naveed, J. Liang, S. Nussberger, The structure of the TOM core complex in the mitochondrial outer membrane. Biological Chemistry 401, 687–697 (2020).

36. D. Gessmann, et al., Structural elements of the mitochondrial preprotein-conducting channel Tom40 dissolved by bioinformatics and mass spectrometry. Biochimica et Biophysica Acta (BBA) - Bioenergetics 1807, 1647–1657 (2011).

37. T. Shiota, et al., Molecular architecture of the active mitochondrial protein gate. Science 349, 1544–1548 (2015).

38. D. Rapaport, W. Neupert, R. Lill, Mitochondrial Protein Import. Journal of Biological Chemistry 272, 18725–18731 (1997).

39. Y. Araiso, K. Imai, T. Endo, Role of the TOM Complex in Protein Import into Mitochondria: Structural Views. Annu. Rev. Biochem. 91, 679–703 (2022).

40. T. Shiota, H. Mabuchi, S. Tanaka-Yamano, K. Yamano, T. Endo, In vivo protein-interaction mapping of a mitochondrial translocator protein Tom22 at work. Proc. Natl. Acad. Sci. U.S.A. 108, 15179–15183 (2011).

41. H. Yamamoto, et al., Dual role of the receptor Tom20 in specificity and efficiency of protein import into mitochondria. Proceedings of the National Academy of Sciences 108, 91–96 (2011).

42. A. C. Y. Fan, et al., Interaction between the Human Mitochondrial Import Receptors Tom20 and Tom70 in Vitro Suggests a Chaperone Displacement Mechanism. Journal of Biological Chemistry 286, 32208–32219 (2011).

43. S. Wang, et al., Spatiotemporal stop-and-go dynamics of the mitochondrial TOM core complex correlates with channel activity. Commun Biol 5, 471 (2022).

44. L. Bolliger, T. Junne, G. Schatz, T. Lithgow, Acidic receptor domains on both sides of the outer membrane mediate translocation of precursor proteins into yeast mitochondria. The EMBO Journal 14, 6318–6326 (1995).

45. N. Morgner, et al., A New Way To Detect Noncovalently Bonded Complexes of Biomolecules from Liquid Micro-Droplets by Laser Mass Spectrometry. Aust. J. Chem. 59, 109–114 (2006).

46. N. Morgner, C. V. Robinson, *Mass* ign: An Assignment Strategy for Maximizing Information from the Mass Spectra of Heterogeneous Protein Assemblies. Anal. Chem. 84, 2939–2948 (2012).

47. E. Gasteiger, et al., ExPASy: The proteomics server for in-depth protein knowledge and analysis. Nucleic Acids Res 31, 3784–3788 (2003).

48. D. Kimanius, L. Dong, G. Sharov, T. Nakane, S. H. W. Scheres, New tools for automated cryo-EM single-particle analysis in RELION-4.0. Biochemical Journal 478, 4169–4185 (2021).

49. S. Q. Zheng, et al., MotionCor2: anisotropic correction of beam-induced motion for improved cryo-electron microscopy. Nat Methods 14, 331–332 (2017).

50. A. Rohou, N. Grigorieff, CTFFIND4: Fast and accurate defocus estimation from electron micrographs. J Struct Biol 192, 216–221 (2015).

51. T. Wagner, et al., SPHIRE-crYOLO is a fast and accurate fully automated particle picker for cryo-EM. Commun Biol 2, 1–13 (2019).

52. A. Punjani, J. L. Rubinstein, D. J. Fleet, M. A. Brubaker, cryoSPARC: algorithms for rapid unsupervised cryo-EM structure determination. Nat Methods 14, 290–296 (2017).

53. E. F. Pettersen, et al., UCSF ChimeraX: Structure visualization for researchers, educators, and developers. Protein Sci 30, 70–82 (2021).

54. R. Evans, et al., Protein complex prediction with AlphaFold-Multimer. 2021.10.04.463034 (2022).

55. P. Emsley, B. Lohkamp, W. G. Scott, K. Cowtan, Features and development of *Coot*. Acta Crystallogr D Biol Crystallogr 66, 486–501 (2010).

56. T. I. Croll, ISOLDE: a physically realistic environment for model building into low-resolution electron-density maps. Acta Cryst D 74, 519–530 (2018).

57. D. Liebschner, et al., Macromolecular structure determination using X-rays, neutrons and electrons: recent developments in Phenix. Acta Cryst D 75, 861–877 (2019).

58. J. Jumper, et al., Highly accurate protein structure prediction with AlphaFold. Nature 596, 583–589 (2021).

