## Supplementary Material for "Two conformations of the Tom20 preprotein receptor in the TOM holo complex"

**This PDF file includes:**

Figures S1 to S11
Tables S1 to S2

### Supplementary Figures and Tables

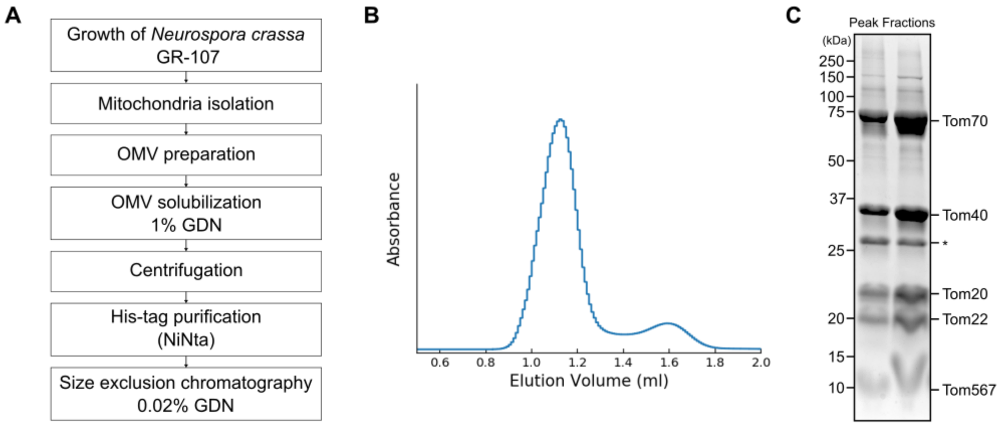

**Fig. S1.**
Purification of the TOM holo complex from outer membrane vesicles. (**A**) Scheme of the TOM holo purification steps. (**B**) Size exclusion chromatography profile of the TOM holo complex in GDN using the Superdex 200 Increase column. (**C**) Coomassie-stained SDS-PAGE of the main peak after size exclusion chromatography. The asterisk indicates VDAC.

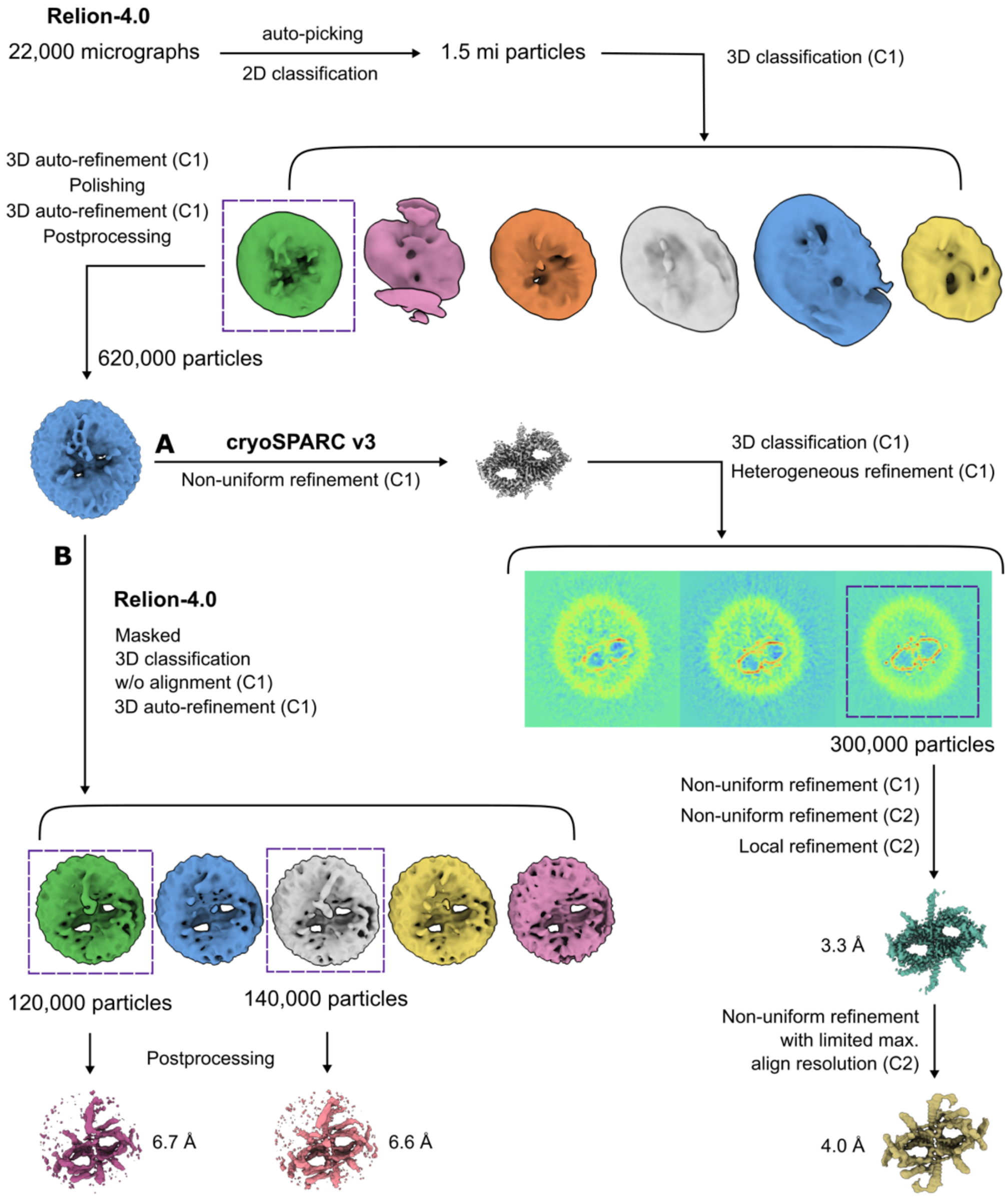

**Fig. S2.**
Single-particle cryoEM analysis of the TOM complex structure. The workflow includes the steps leading to: (**A**) The 3.3 Å resolution map of the TOM core complex and the 4 Å resolution map of the presequence bound TOM core complex in cryoSPARC v3. (**B**) The TOM core + Tom20 complex maps in conformation $C_1$ at 6.7 Å and $C_2$ at 6.6 Å resolution.

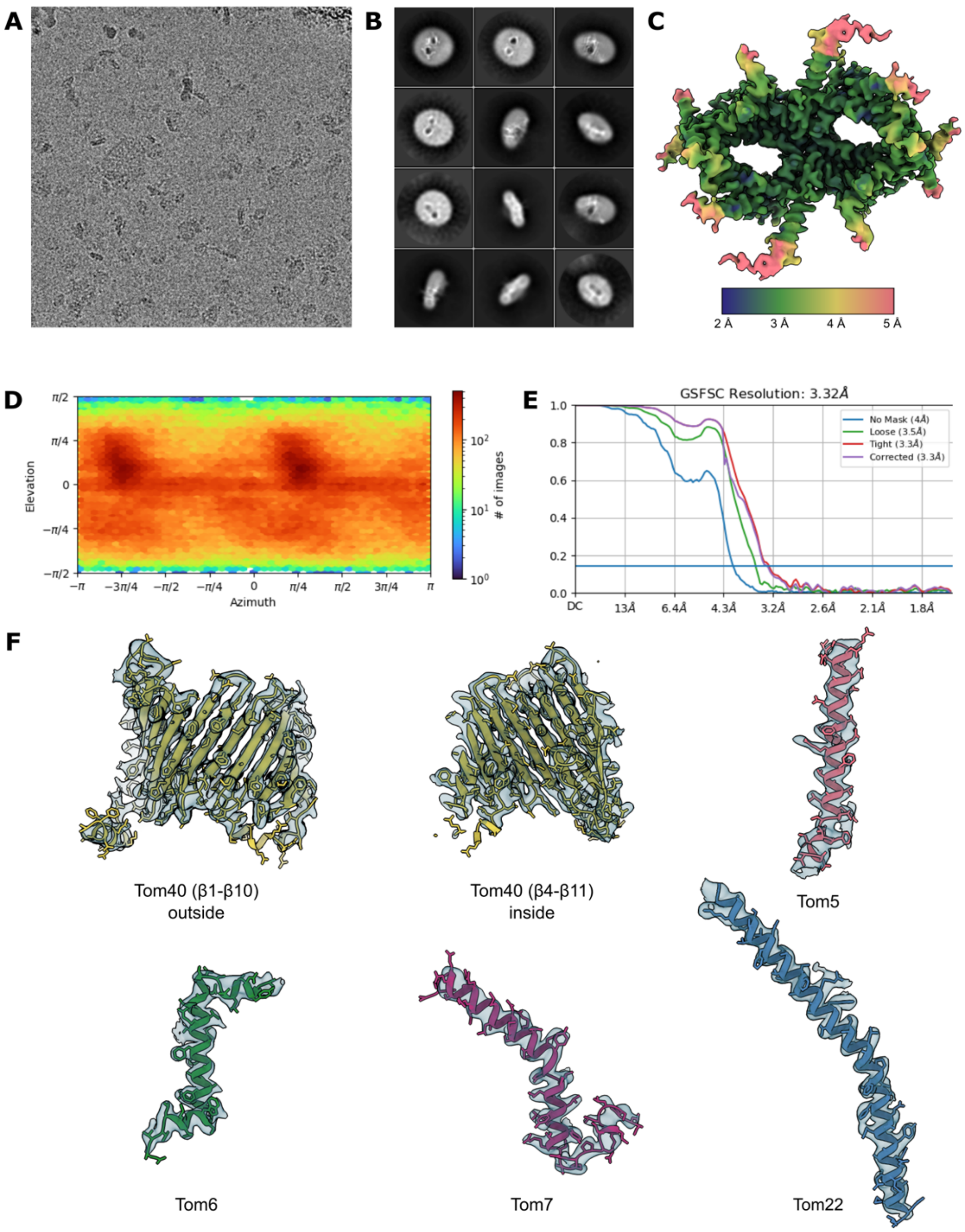

**Fig. S3.**
Single-particle processing of the TOM core complex. (**A**) Representative electron micrograph of the cryoEM collection. (**B**) Representative 2D class averages of the TOM complex. (C) Final reconstructed map colored according to local resolution as estimated by cryoSPARC. (**D**) Particle distribution in the final reconstruction presented as a heat map as measured in cryoSPARC. (**E**) Fourier shell correlation of final local refinement and local resolution estimation carried out in cryoSPARC. (**F**) The individual subunits of the TOM core complex display model quality.

A

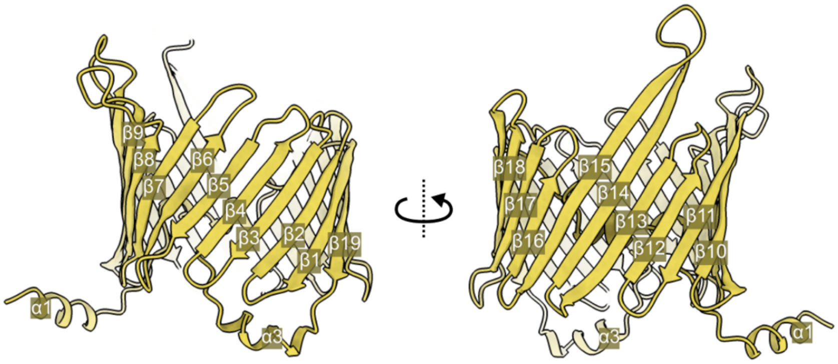

B

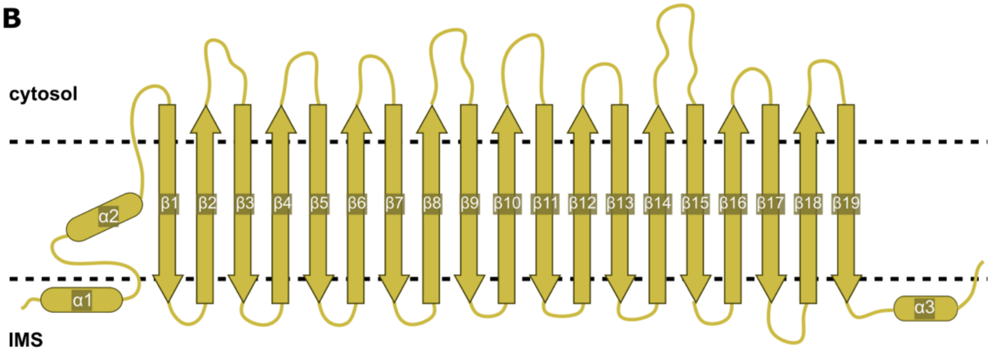

**Fig. S4.**
The Tom40 translocation pore. (**A**) Atomic model of Tom40 with numbered ß-strands. (**B**) Schematic diagram of Tom40 secondary structure.

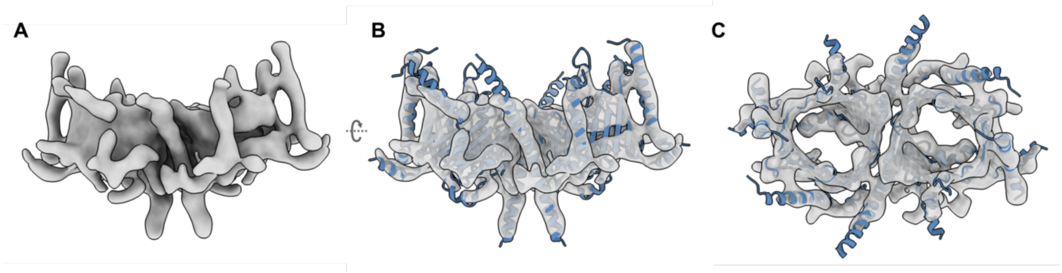

**Fig. S5.**
Superposition of the NcTOM core model (PDB 8B4I, blue), derived from our present 3.32 Å map, with on our previous 6.8 Å map (EMDB-3761) (**A**), as seen from the side (**B**) and from the cytosol (**C**).

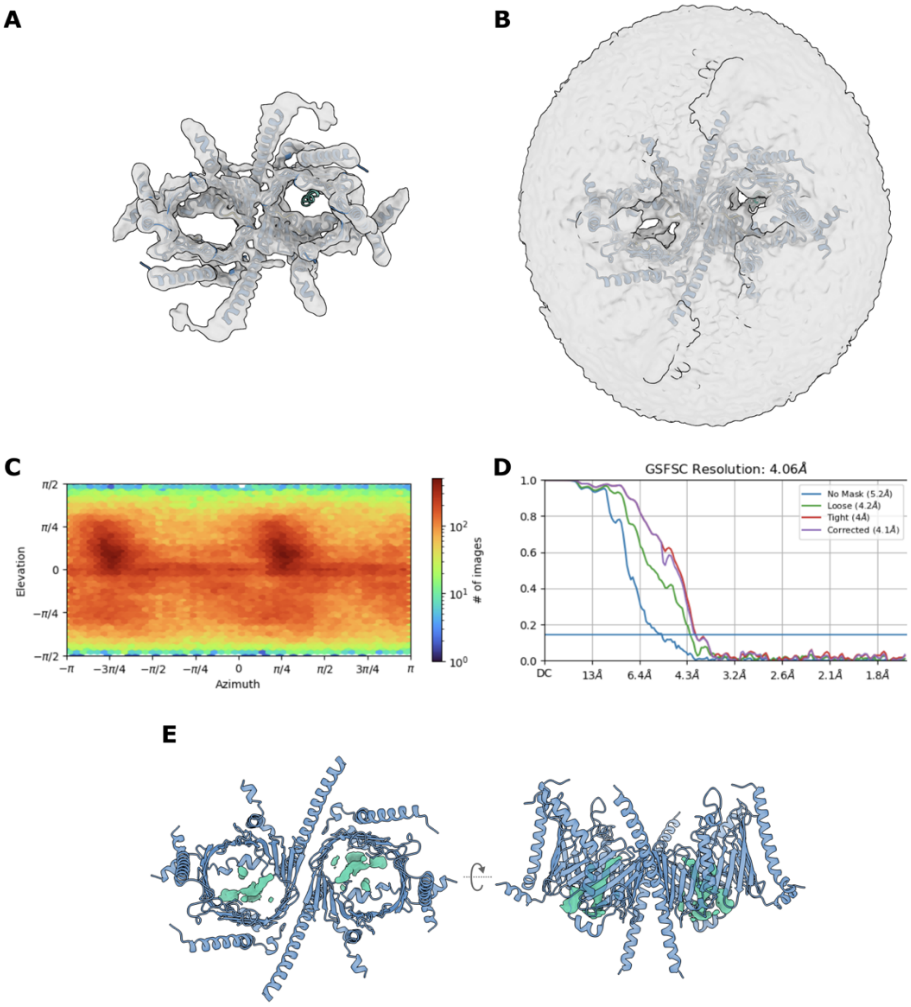

**Fig. S6.**
Processing results of the TOM core complex with bound presequence. Map obtained by non-uniform refinement with limited alignment resolution of the TOM core particles with C2 symmetry applied. Superposition of the 4 Å map with the TOM core model and the rigid-body-fitted pALDH inside one pore at (**A**) high and (**B**) low density threshold. (**C**) Particle distribution in the final reconstruction presented as a heat map measured in cryoSPARC. (**D**) Fourier shell correlation of final local refinement and local resolution estimate from cryoSPARC. (**E**) Difference map showing the presequence density superposed on the TOM core model as seen from the cytosol (left) or from the membrane.

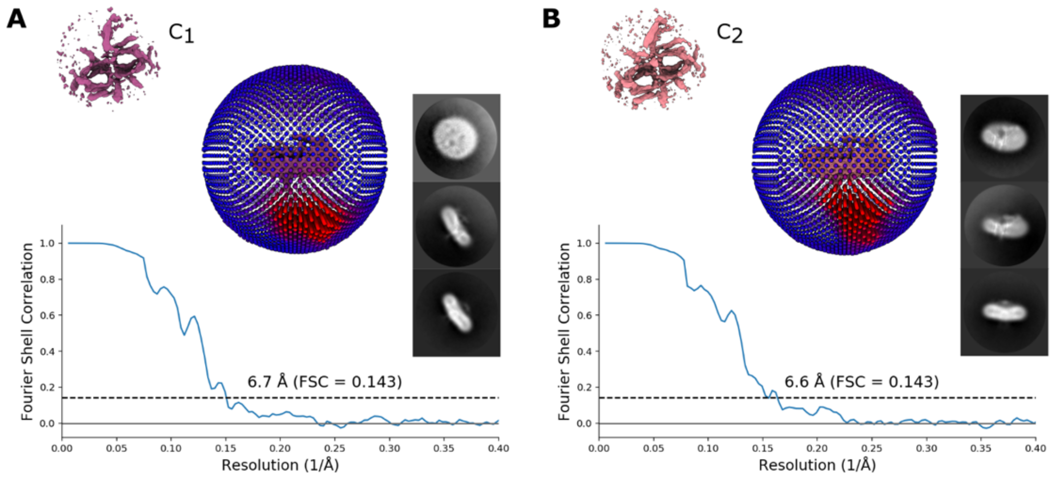

**Fig. S7.**
Image processing of the NcTOM core + Tom20 complex in conformations (**A**) $C_1$ and (**B**) $C_2$. Each figure shows representative 2D averages of the final particles in each map, as well as the particle distribution in the final reconstruction and the Fourier shell correlation of the final refinement measured in Relion-4.0.

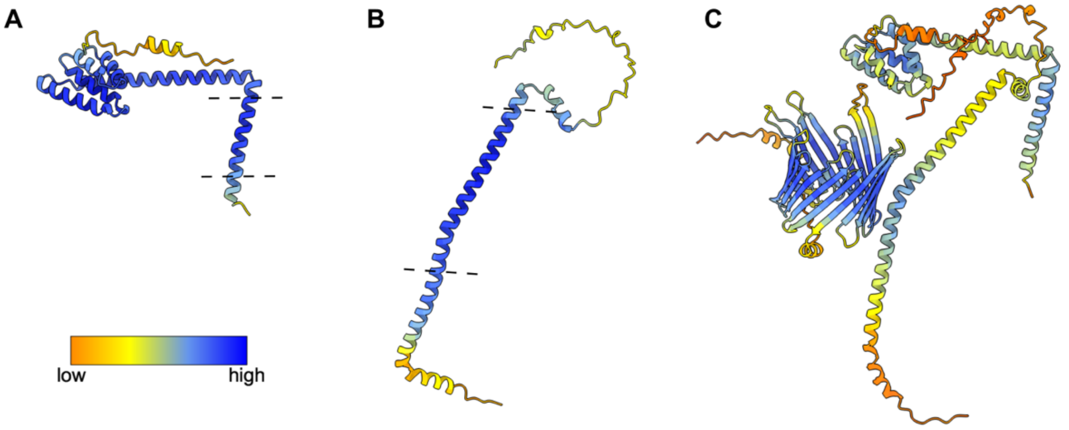

**Fig. S8.**
AlphaFold predictions of key TOM subunits colored by estimated per-residue confidence. Orange indicates low confidence and blue high confidence. (**A**) and (**B**) show the monomer predictions of Tom20 and Tom22 that were used for rigid-body-fitting. Dashed lines indicate the likely position of the outer mitochondrial membrane, based on the hydrophobicity of modelled transmembrane helices. (**C**) AlphaFold-Multimer prediction model of the $Tom20_1Tom22_1Tom40_1$ subcomplex.

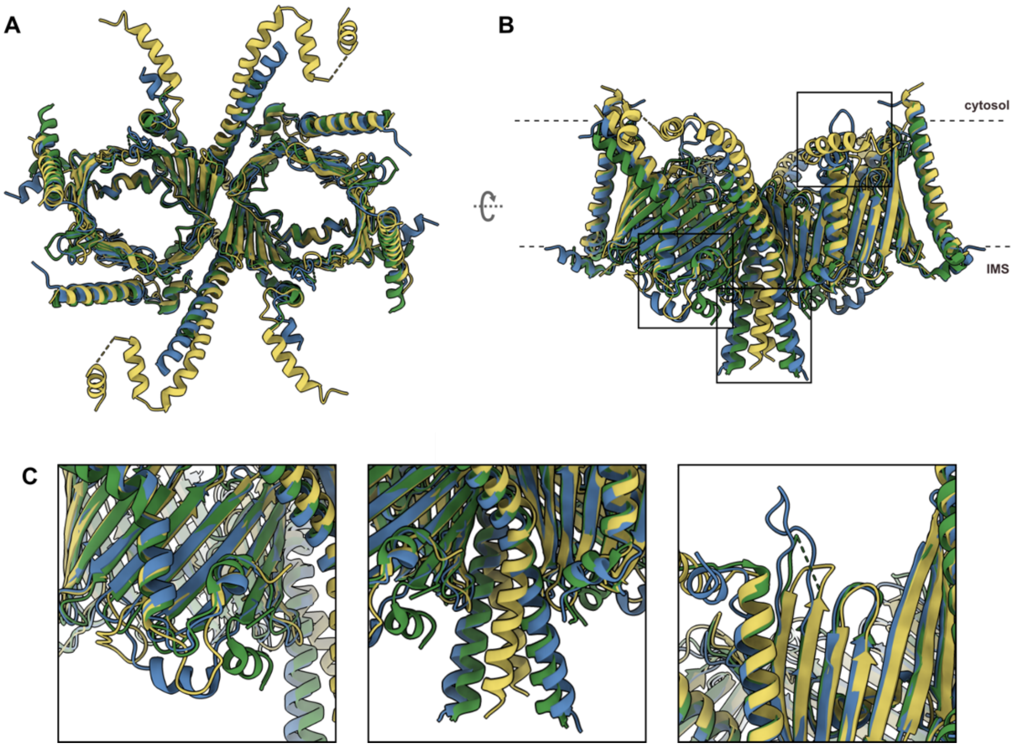

**Fig. S9.**
Differences between the models of the TOM core complex from human, yeast and *N. crassa* mitochondria. Human TOM is shown in yellow (PDB 7CP9), yeast TOM in green (PDB 6UCU) and *N. crassa* TOM in blue (PDB 8B4I). (**A**) Cytosolic view of the three models. (**B**) Side view of the models with squares highlighting areas of interest. Dashed lines indicate the outer membrane. (**C**) Close-up of differences in Tom7, Tom22 and Tom40.

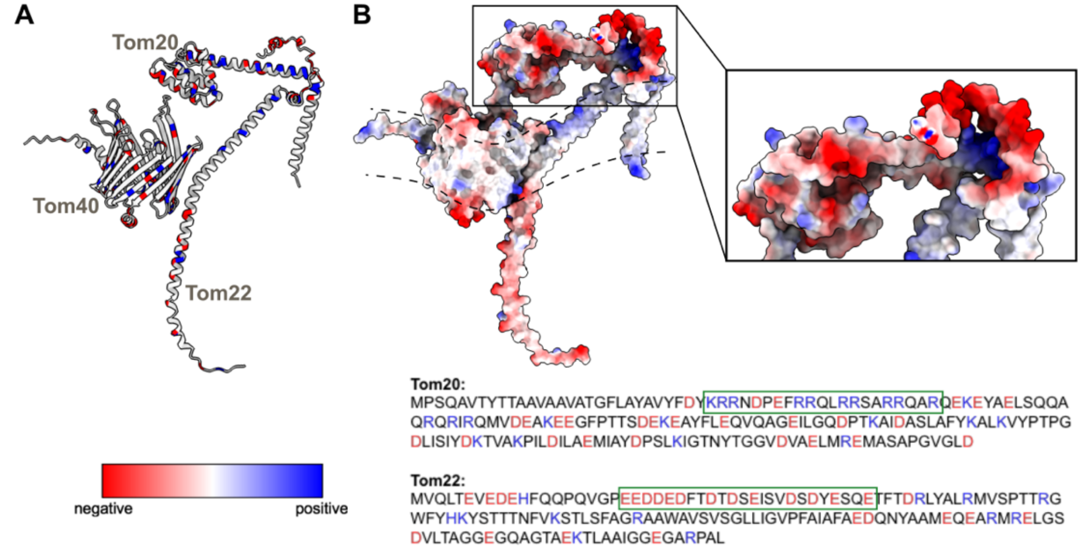

**Fig. S10.**
AlphaFold prediction of the $Tom20_1Tom22_1Tom40_1$ subcomplex colored by electrostatic potential. Red and blue indicate negatively and positively charged regions, respectively. (**A**) Cartoon representation with highlighted charged regions. (**B**) Close-up of the predicted docking site of Tom20 on Tom22. The N-Terminal of Tom22 is cut transversally to show the region of interest. The likely position of the outer membrane is indicated by dashed lines. Sequences of Tom20 and Tom22 are shown, color coded by residue charge. Green boxes highlight the regions of charge complementarity in Tom20 and Tom22 that we propose holds the two subunits together by electrostatic interactions.

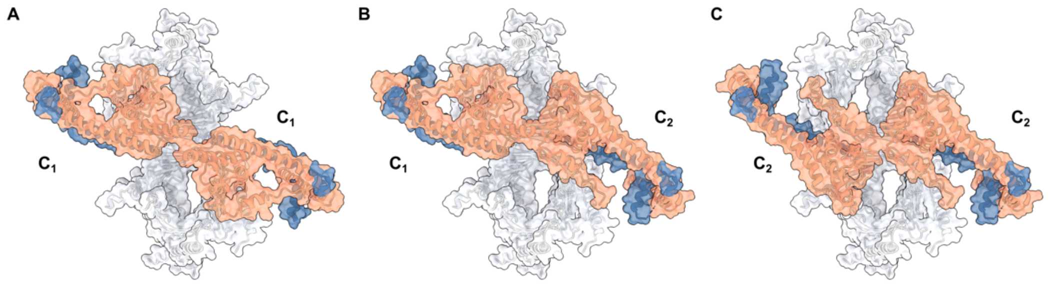

**Fig. S11.**
We attempted to fit two copies of Tom20 as rigid bodies into our TOM core dimer model, in its two different conformations: (**A**) $C_1$ + $C_1$, (**B**) $C_1$ + $C_2$ and (**C**) $C_2$ + $C_2$. The receptor domains of our fitted Tom20 clash in the cytosol in both conformations, making it unlikely that they can simultaneously coexist.

**Table S1.** Predicted and experimental mass related to the TOM holo complex, subcomplexes and subunits. The predicted masses were calculated using the ExPASy tool. The value for the unidentified sT = 5.975 kD was taken from the average of the mass of Tom5, Tom6 and Tom7.

| Protein | Predicted Mass (kDa) | LILBID Mass (kDa) |
|---|---|---|
| Tom5 | 5.402 | 5.657 |
| Tom7 | 6.061 | 6.231 |
| Tom6 | 6.463 | 6.550 |
| Tom22-6His | 17.639 | 17.901 |
| Tom20 | 20.228 | 20.099 |
| Tom40 | 38.150 | 37.925 |
| Tom70 | 69.340 | 69.363 |
| $Tom20_2$ | 40.456 | 39.785 |
| $sT_1Tom40_1$ | 44.125 | 44.020 |
| $sT_2Tom40_1$ | 50.101 | 53.483 |
| $Tom22_1Tom40_1$ | 54.966 | 55.553 |
| $Tom20_2Tom22_1$ | 57.272 | 57.623 |
| $sT_1Tom22_1Tom40_1$ | 60.941 | 62.650 |
| $sT_2Tom22_1Tom40_1$ | 66.917 | 68.859 |
| $Tom20_1Tom22_1Tom40_1$ | 75.194 | 75.692 |
| $sT_2Tom70_1$ | 81.291 | 81.980 |
| $sT_1Tom20_1Tom22_1Tom40_1$ | 81.169 | 82.857 |
| $Tom22_1Tom70_1$ | 86.156 | 86.453 |
| $sT_2Tom20_1Tom22_1Tom40_1$ | 87.145 | 87.762 |
| $Tom20_1Tom70_1$ | 89.568 | 89.496 |
| $Tom40_1Tom70_1$ | 107.490 | 107.005 |

**Table S2.** CryoEM data collection, refinement and validation of the NcTOM core complex.

| | **TOM core** |
|---|---|
| **Data collection and processing** | |
| Magnification | 105kx |
| Voltage (kV) | 300 |
| Electron exposure | 55 e-/A2 |
| Defocus Range (μm) | -1.2 to - 3.0 |
| Pixel size (Å) | 0.837 |
| Symmetry imposed | C2 |
| Initial particles | 1,499,000 |
| Final particles | 304,506 |
| Map resolution (Å) | 3.32 |
| FSC Threshold | 0.143 |
| **Refinement** | |
| Initial model used | AlphaFold-Multimer |
| Model resolution | 3.24 |
| FSC Threshold | 0.143 |
| Map sharpening B factor (Å2) | -80 |
| **Model composition** | |
| Nonhydrogen atoms | 8,250 |
| Protein residues | 1,026 |
| Ligands | 9 |
| **R. m. s. deviations** | |
| Bond lengths (Å) | 0.004 |
| Bond angles (˚) | 0.571 |
| **Validation** | |
| MolProbity score | 1.17 |
| Clashscore | 3.81 |
| Poor rotamers (%) | 0.00 |
| Ramachandran plot | |
| Favored (%) | 98.51 |
| Allowed (%) | 1.49 |
| Disallowed (%) | 0.00 |
